# Inhibition of broomrape germination by 2,4-diacetylphloroglucinol produced by environmental *Pseudomonas*

**DOI:** 10.1101/2023.03.01.529533

**Authors:** Tristan Lurthy, Ségolène Perot, Florence Gerin-Eveillard, Marjolaine Rey, Florence Wisniewski-Dyé, Jordan Vacheron, Claire Prigent-Combaret

**Affiliations:** Ecologie Microbienne, Université Claude Bernard Lyon1, Université de Lyon, CNRS UMR-5557, INRAe UMR-1418, VetAgro Sup, 43 Boulevard du 11 Novembre 1918, 69622 Villeurbanne, France; Department of Fundamental Microbiology, University of Lausanne, Lausanne, Switzerland

**Keywords:** Environmental *Pseudomonas*, 2,4-diacetylphloroglucinol, DAPG, broomrape *Phelipanche* sp., *Orobanche* sp., parasitic weeds

## Abstract

Phloroglucinol compounds (PGCs) produced by environmental *Pseudomonas* are well known for their capacity to limit plant-pathogen infection. Although PGCs and more specifically 2,4-diacetylphloroglucinol (DAPG) are well studied for their antimicrobial properties, they are to some extent toxic for crop plants. Parasitic weeds such as broomrapes (*Phelipanche ramosa* and *Orobanche cumana*) cause severe damage to crops and their development must be controlled. Here, we assessed the potential herbicidal effect of the bacterial model *Pseudomonas ogarae* F113, a PGCs-producing bacterium, on parasitic weed germination. We show using a mutagenesis approach that PGCs produced in bacterial supernatants are the main determinant inhibiting the germination of broomrapes. The use of individual or cocktails of pure PGCs revealed that the inhibition of the germination depends on the PGCs molecular structure and their concentrations as well as the broomrape species and pathovars. Furthermore, the inhibition caused by the PGCs is irreversible, causing a brown coloration of the broomrape seeds. Then, we evaluated in non-sterile soils the ability of bacterial inoculants or chemical DAPG to limit the infection of broomrapes on oil seed rape. Only the inoculation of PGCs-producing bacteria limited the infection of *P. ramosa*. Moreover, elemental profiling analysis of oil seed rape revealed that neither the inoculant nor applied DAPG affected the nutrition capacity of the oil seed rape. Our study expands the knowledge on the role that these multi-talented plant-beneficial *Pseudomonas* play in the environment and open new avenues for the development of natural bioherbicides to ward off parasitic plant infection.

## Introduction

Broomrapes are parasitic plants causing significant damage to different crops in agroecosystems [1]. They belong to the *Orobanche* and *Phelipanche* genera from the *Oroabanchaceae* family [2, 3]. These parasitic plants are obligate root holoparasites, entirely dependent on their host plant to survive as they are not capable of photosynthesis [4]. Indeed, these parasitic plants obtain all the resources they need while maintaining their host alive for accomplishing their entire life cycle [5, 6]. The seed germination and haustorium formation are specifically induced by allelochemical signals such as strigolactones released from the host roots [7]. The activity spectrum of these parasitic plants can be either specific (e.g., *Orobanche cumana* which can only parasitize sunflower) or generalist (e.g., *Phelipanche ramosa* able to parasitize hemp (*Cannabis sativa*), tobacco (*Nicotiana tabacum*), and oilseed rape (*Brassica napus*)) [1, 5]. Broomrapes produce a large number of small seeds (less than 3 mm) that can survive in soil for several decades. This constitute the main problem to constrain their deleterious impact on crops [8]. The survival of the seeds depends on various abiotic factors (pH, humidity, climate) [9] and biotic factors (host plants, soil and rhizosphere microbiota; [6, 10–12]. Different agricultural strategies attempt to regulate broomrape populations in agroecosystems, such as crop rotation, triggering the suicidal-germination of the plant parasitic seeds or the use of resistant host plant varieties or chemical herbicides [5]. However, biological control solutions are emerging to limit broomrape infestation, including the use of microorganisms [5, 13]. Indeed, several microorganisms inhibit the germination of different broomrape species, including, among others, *Fusarium oxysporum* [14], *Azospirillum brasilense* [15] or *Pseudomonas fluorescens* [16]. Although the use of microorganims represents a promising alternative to ward off parasitic plants, their mode of action as well as the identification of the metabolites responsible of their inhibition effect remains often uncharted.

Several environmental *Pseudomonas* are well known as plant-colonizing bacteria [17]. These *Pseudomonas* usually display a large arsenal of secondary metabolites encoded within their genomes that act on plant health and development [18, 19]. Among these metabolites, 2,4-diacetylphloroglucinol (DAPG) has been studied notably for its role in plant protection. DAPG and its biosynthetic intermediates, phloroglucinol (PG) and monoacetylphloroglucinol (MAPG) are the main phloroglucinol compounds (PGCs) produced by *Pseudomonas* belonging to the *P. protegens* and *P. corrugata* subgroups [20]. The production of DAPG relies on the presence of the *phl* gene cluster composed out of nine genes [21, 22]. The initiation of the synthesis of PG from malonyl-CoA is mediated by *phlD* encoding a polyketide synthase, while *phlABC* encode for enzymes implicated into the transformation of PG to MAPG and subsequently to DAPG. The transformation of MAPG to DAPG is reversible through a hydrolase encoded by *phlG*. The remaining genes, *phlF/phlH* and *phlE*, are involved in the regulation as well as the secretion of these PGCs, respectively. The production of PGCs by *Pseudomonas* is influenced by environmental factors including carbon sources [23] or specific metabolites found in the root exudates such as flavonoids, apigenin and phloretin [24].

In addition to having been studied for its role in plant pathogen suppression, DAPG acts as a signal molecule affecting gene expression of plant-beneficial traits in other microorganisms. Indeed, DAPG was described as an inducer of the production of PGCs and a repressor the production of pyoluteorin in other *Pseudomonas* [25, 26]. Moreover, DAPG produced by *Pseudomonas* also activates the expression of genes involved in the production of auxins by *Azospirillum baldaniorum* Sp245, another plant-beneficial microorganism [27]. Since PGC-producing *Pseudomonas* are residing in the vicinity of or on plant roots, PGCs produced diffuse and also interact directly with plant root cells. Thus, it was shown that DAPG elicited the plant induced systemic resistance (ISR), protecting partially the plant leaves from the oomycete *Peronospora parasitica* [28, 29]. On the root part, the addition of DAPG triggered a massive increase of the efflux of amino acids by plant root cells [30]. It was also demonstrated that DAPG modulates auxin-dependent plant signaling pathway leading to significant modifications of plant root development [31–33]. Moreover, following an exposition of DAPG, the germination as well as the development of different crop plants were severely impacted [31, 34, 35]. Nevertheless, this herbicidal effect remains variable according to the plant species and was observed following the exposure of high concentrations that do not reflect those produced *in vivo*.

In this study, we aimed to investigate the impact of PGCs on the germination of the two main parasitic plants, *Phelipanche* and *Orobanche*. To evaluate the herbicidal effects of these PGCs, we have investigated the inhibitory effect of PGCs-producing strains and pure molecules in different *in vitro* and *in planta* experimental systems. First, the impact of the PGCs producing strain *Pseudomonas ogarae* F113 and its mutants, a PGC-deficient and PGC overproducers, have been studied on broomrape germination by applying culture supernatants. Then, the role of PGCs on the germination of four different broomrapes was assessed at different concentrations under *in vitro* experiment. Finally, we evaluated the ability of PGCs-producing bacteria and DAPG application to protect oilseed rape against broomrape in greenhouse.

## Materials and methods

### Bacterial strains and media

We used the plant-beneficial model strain *Pseudomonas ogarae* F113 (formerly named *P. fluorescens* F113 and *P. kilonensis* F113) [23] and several of its mutants [33]. The bacterial strains used in this study as well as their characteristics are listed in **Supplementary Table 1**. The different bacterial strains were incubated at 28°C in King’s B [36] medium or in a modified AB medium (ABm) supplemented with gentamycin (15 μg.mL^-1^ when necessary) to maintain plasmid pBBR1-MCS5-*phlD*. ABm was composed of salts [MgSO_4_ (1.2 mM), CaCl_2_ (70 μM), NH4Cl (18 mM), KCl (2 mM), FeSO_4_ (9 μM)], a phosphate buffer diluted ten-fold containing K_2_HPO_4_ (1.725 mM) and NaH_2_PO_4_ (960 μM) and fructose (20 mM) as carbon source.

### Plant material

Seeds of *Phelipanche ramosa* were collected in France as described in Huet et al. 2020 on winter oilseed rape (*Phelipanche ramosa* pv. oilseed rape), tobacco (*Phelipanche ramosa* pv. tobacco) and hemp (*Phelipanche ramosa* pv. hemp). Seeds of *Orobanche cumana* that parazites sunflower were provided by Terres Inovia in 2016. Seeds of *Brassica napus* cultivar AMAZZONITE (broomrape-sensitive) were provided by the breeder companies RAGT 2n (France).

### Chemicals

The germination of broomrape seeds was triggered using the synthetic strigolactone analogue GR24 (Chiralix, Nijmegen, NL). It was first suspended in acetone (4.79 mg.mL^-1^), then diluted at 10 μM with a phosphate buffer (1 mM sodium-potassium phosphate buffer at pH 7.5).

Phloroglucinol (PG, Sigma-Aldrich), mono-acetyl-phloroglucinol (MAPG, Cayman Chemical), 2,4-diacetyl-phloroglucinol (DAPG, ChemCruz) and tri-acetyl-phloroglucinol (TAPG, Santa Cruz Biotechnology) were suspended in methanol (20 mM). These solutions were diluted with 1 mM phosphate buffer to obtain different stock solutions at different concentrations (66.60, 33.30, 16.65, 8.33, 4.16 μM). As methanol might have an effect on the germination of broomrapes, the final concentration was adjusted to 0.33% in all these stock solutions to prevent an effect of the dilution.

### Quantification of phloroglucinol compounds produced in bacterial supernatants

The quantification of PGCs was conducted on 1.5 mL of bacterial culture supernatants for each condition. First, supernatants were lyophilized (Martin Christ Alpha 1-4 LSC, Osterode, Germany) prior solid/liquid extraction with methanol. Samples were sonicated 20 min, then centrifuged for 20 minutes at 15 000 *g* and the supernatants recovered. The extraction protocol was repeated, leading to a total extracted volume of 3 mL per sample. The organic phase (methanol) was dried using a SpeedVac (Centrivap Cold Trap Concentrator; LABCONCO Co., MO, USA). Dried extracts were suspended in 200 μL of methanol and centrifuged for 5 min at 12 000 g to pellet the remaining solid phase.

Two hundred microliters of the supernatant were then transferred into vials and were proceeded for ultra-high pressure liquid chromatography coupled with UV (UHPLC-UV) analysis, as described in Rieusset et al. [42]. Chromatograms were analysed with MassHunter Qualitative Analysis B.07.00 software (Agilent Technologies®) and the quantification of DAPG and other PGCs was done according to a standard curve with commercial PGCs.

### Inhibition of broomrape germination *in vitro*

Broomrape seeds were surface-disinfected according to [37] with minor modifications. Briefly, broomrape seeds were soaked 5 min in a bleach solution (9.6% active chlorine) and then washed 5 times with sterile water. After washing, 1 mM phosphate buffer supplemented with plant agar 0.1% and PPM 0.2% (Plant Preservative Mixture; Plant Cell Technology) was added to obtain a density of approximatively 2000 seeds.mL^-1^. These solutions containing the seeds were conditioned in sealed tubes for 10 days at 21°C in the dark in a cooled incubator (LMS, model 120, Kent, UK). The supernatant of conditioned seeds was removed and replaced by fresh phosphate buffer supplemented with plant agar 0.1% and PPM 0.2%. Fifteen microliters of this seed suspension were distributed in a 96-well plate (Cellstar®; Greiner Bio-One, France), corresponding to approximatively 30 seeds per well. Then, 10 μL of GR24 solution were then added in each well (final concentration of 1 μM). Seventy-five microliters of either bacterial supernatants (3-fold diluted) or cocktails or individual PGCs were added to obtain a final volume of 100 μL per well. For the addition of PGCs, the 75 μL were taken from the different stock solutions described above to obtain final concentrations of 50, 25, 12.5, 6.25 and 3.125 μM. Negative controls were realized using 75 μL of fresh ABm medium fructose 20 mM for supernatant (3-fold diluted) or phosphate buffer with 0.33% of methanol. After 10 days of incubation at 21°C in the dark, the percentage of broomrape germinated seeds was counted under a binocular (Leica, Switzerland) using the software Zen 2.3.

### Greenhouse experiments

*Brassica napus* plants were grown on a soil mix containing 1/3 of a natural loamy soil collected at the experimental farm in La Côte-St-André (France; 16.2% clay, 43.9% silt and 39.9% sand, pH 7.0, in water; 2.1% organic matter [38], 1/3 of vermiculite and 1/3 of TS3 peat-based substrate (Klasmann-Deilmann GmbH, Geeste, Germany). The humidity of the soil mix was maintained at 70% of field capacity. Each pot was filled with 1 liter of free-broomrape soil mix and then further filled with another liter of soil mix contaminated with non-disinfected seeds of *Phelipanche ramosa* pv. oilseed rape leading to a final density of 3.9 mg of seeds per pot corresponding to approximatively 300 seeds per liter of soil. Seeds of *Brassica napus* cultivar AMAZZONITE were sown in pot after being pre-germinated 24h in the dark at 21°C in Petri dish containing water-soaked Whatman paper.

Two experiments were performed under greenhouse conditions. The first one was the inoculation of F113 and its mutant impaired in the production of PGCs (Δ*phlD*). The different bacterial strains were cultured 24h in King’s B medium at 28°C. The bacteria were centrifuged at 4500 rpm during 10 min and washed with a MgSO_4_ 10 mM solution before being adjusted to a bacterial concentration of 2.10^6^ CFU.mL^-1^. Five milliliters of these bacterial suspensions were sprayed at the base of the plant stem. Five milliliters of a MgSO_4_ 10 mM solution were applied as non-inoculated control. The second experiment was the application of DAPG. Five milliliters of a solution concentrated at 50 μM or 250 μM were applied respecting the same amount of MeOH solvent of 1.25% (e.g. 10 mL of solution DAPG 250 μM = 125 μL solution DAPG 20mM + 9.875 mL of 1 mM phosphate buffer pH 7.5). The control condition without DAPG corresponds to the application of 5 mL of 1 mM phosphate buffer pH 7.5 containing 1.25% of methanol.

Twenty pots per conditions were used for the inoculation of bacteria while 10 pots per conditions were used for the DAPG experiment. The shoot and root dry biomasses of *B. napus* were measured at the end of the experiment after 3 days at 70 °C in an oven. The experiment was conducted under controlled conditions with a 16h light and 8h dark photoperiod, at 25°C with 50-70% relative humidity in a greenhouse for 50 days. Treatments were applied twice during the experiment. The first application was performed when *B. napus* had between 2 to 4 leaves. The second application was done at 6 to 8 leaves.

### Elemental profiling analysis of *B. napus*

After 50 days, dry samples from shoot of rapeseed from greenhouse experiment were ground into fine powder using a metal ball in a 50 mL plastic tube. For the elemental profiling analysis, 5 independent samples of 4 plants for strains-inoculation and of 2 plants for DAPG-treatment were collected. The concentrations of 17 elements (Na, Mo, Cd, Be, B, Mg, P, S, Ca, Mn, Fe, Co, Ni, Cu, Zn and K) were measured by High Resolution Inductively Coupled Plasma Mass Spectrometry (HR ICP-MS, Thermo Scientific, Element 2TM, Bremen, Germany) as described in Lurthy et al. [39].

### Data processing and statistical analysis

Data were analysed using R studio (v.4.2.1) and considered significantly different when p-value < 0.05. The data were assessed for Normal distribution and variance homogeneity using Shapiro-Wilk tests and Bartlett tests respectively. When these parameters were respected, we performed ANOVA coupled with either HSD-Tukey test (more than 3 conditions to compare) or LSD-Fisher test (3 conditions to compare). Otherwise, Kruskal-Wallis tests applying Bonferroni correction were used to detect differences between conditions. A two-way ANOVA was performed to assess the effect of the different variable tested, such as the effect of the pathovar and / or the PGCs applied on broomrape seed germination.

## Results

### The DAPG produced by *Pseudomonas* contributes to the inhibition of the germination of *Phelipanche ramosa in vitro*

We tested the ability of *Pseudomonas ogarae* F113, a PGC-producing *Pseudomonas*, to inhibit the germination of four different broomrapes selected according to their host specificity, their parasitism cycle and their associated microbiota [10]: *P. ramosa* pv. oilseed rape*, O. cumana* sunflower, *P. ramosa* pv. tobacco and *P. ramosa* pv. hemp.

We first determined the concentration of four different PGCs (PG, MAPG, DAPG and TAPG, **Figure 1B**) by UHPLC-UV in the supernatant of *P. ogarae* F113 wild type (F113) as well as in the supernatants of a mutant impaired in the production of DAPG (Δ*phlD*), a complemented mutant (Δ*phlD* Comp.) and a strain engineered to overexpress the gene *phlD* (Over *phlD*) involved in the production of PG, the precursor of the DAPG (**Figure 1A**). Three out of the four different PGCs measured (PG, MAPG and DAPG) were detected in all supernatants, except in the supernatant of Δ*phlD* (**Figure 1B**). The concentration of DAPG in the supernatant of the complemented and the overproducing strains were in the same order of magnitude than F113. However, Δ*phlD* Comp. and Over *phlD* accumulated between 10 to 20 times more PG and MAPG in their supernatants than F113. Finally, TAPG was not detected in any of the bacterial supernatants (**Figure 1B**).

**Figure 1:**
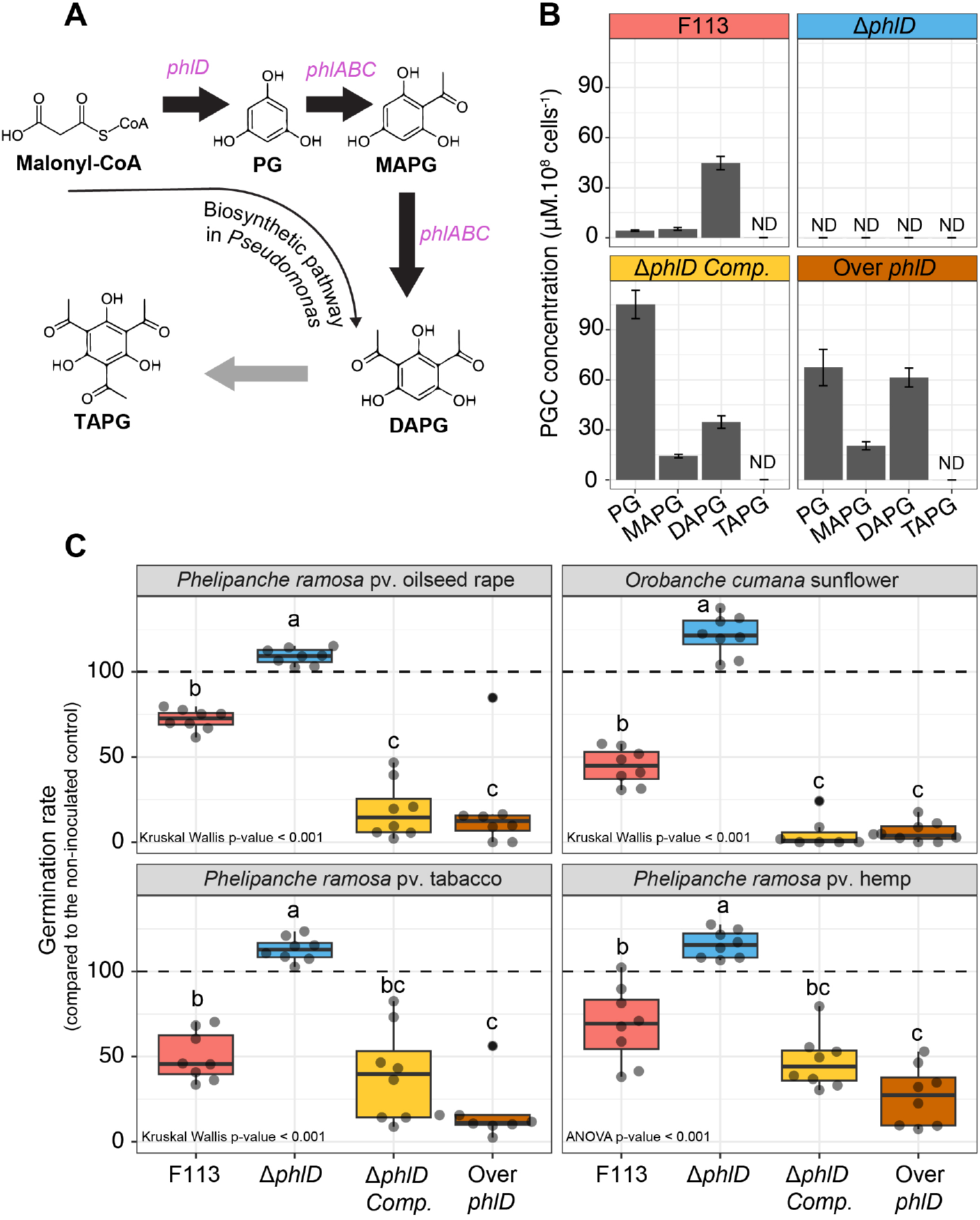
Impact of bacterial supernatants of PGC-producing *Pseudomonas* on the germination rate of different broomrapes. **A:** Biosynthetic pathway of the DAPG in *Pseudomonas* cells (according to Biessy and Filion, 2021). The genes involved in the different transformation steps are written in italic. **B**: Quantification of Phloroglucinol (PG), monoacetylphloroglucinol (MAPG), diacetylphloroglucinol (DAPG) and triacetylphloroglucinol (TAPG) in the bacterial supernatants of *Pseudomonas ogarae* F113 and its mutant derivatives: Δ*phlD* (mutant impaired in the production of DAPG); Δ*phlD* Comp. (complemented mutant strain overproducing PG) and Over Δ*phlD* (DAPG and PG overproducing strain). Error bars correspond to the standard deviation; ND: Not detected. **C**: Impact of these supernatants on the germination capacity of different *P. ramosa* pathovars and *O. cumana in vitro*. The supernatants as well as the control condition were supplemented with 1 μM of the germination stimulant (GR24). Results are expressed as percentage of germination of the non-inoculated ABm medium control. Statistical differences were assessed by ANOVA and Kruskal-Wallis test using a Bonferroni correction and are indicated with letters. The horizontal lines indicate the interquartile range with the center representing the median.

Then, we tested the capacity of the different bacterial supernatants to inhibit the germination of the different broomrapes (**Figure 1C**). We observed that the supernatant of F113 reduced the germination rate of all broomrapes tested (from 28% to 55%). However, the supernatant of Δ*phlD* did not inhibit the germination rate and appears, on the contrary, to slightly promote it compared to the control (>100%) (**Figure 1C**). The highest inhibition of the germination was observed with the supernatants of the complemented and the overproducing strains (**Figure 1C**). Furthermore, the sensitivity to the bacterial supernatants is different according to the broomrape tested (e.g., *P. ramosa* pv. tobacco being more-inhibited by the supernatant of F113 than *P. ramosa* pv. oil seed rape) (**Supplementary Figure 1**).

### Each PGC contributes differentially to the inhibition of broomrape germination

To assess the contribution of the PGCs detected in bacterial supernatants on broomrape germination, we composed three different cocktails made of commercially available PGCs, mimicking the proportions detected in the supernatants of the different F113 derivatives (**Figure 2A**). The application of these cocktails on seeds of *P. ramosa* pv. oilseed rape allowed us to determine whether the effect observed with the complex supernatants was mainly due to the presence of these PGCs and not to other compounds produced by the bacteria.

**Figure 2:**
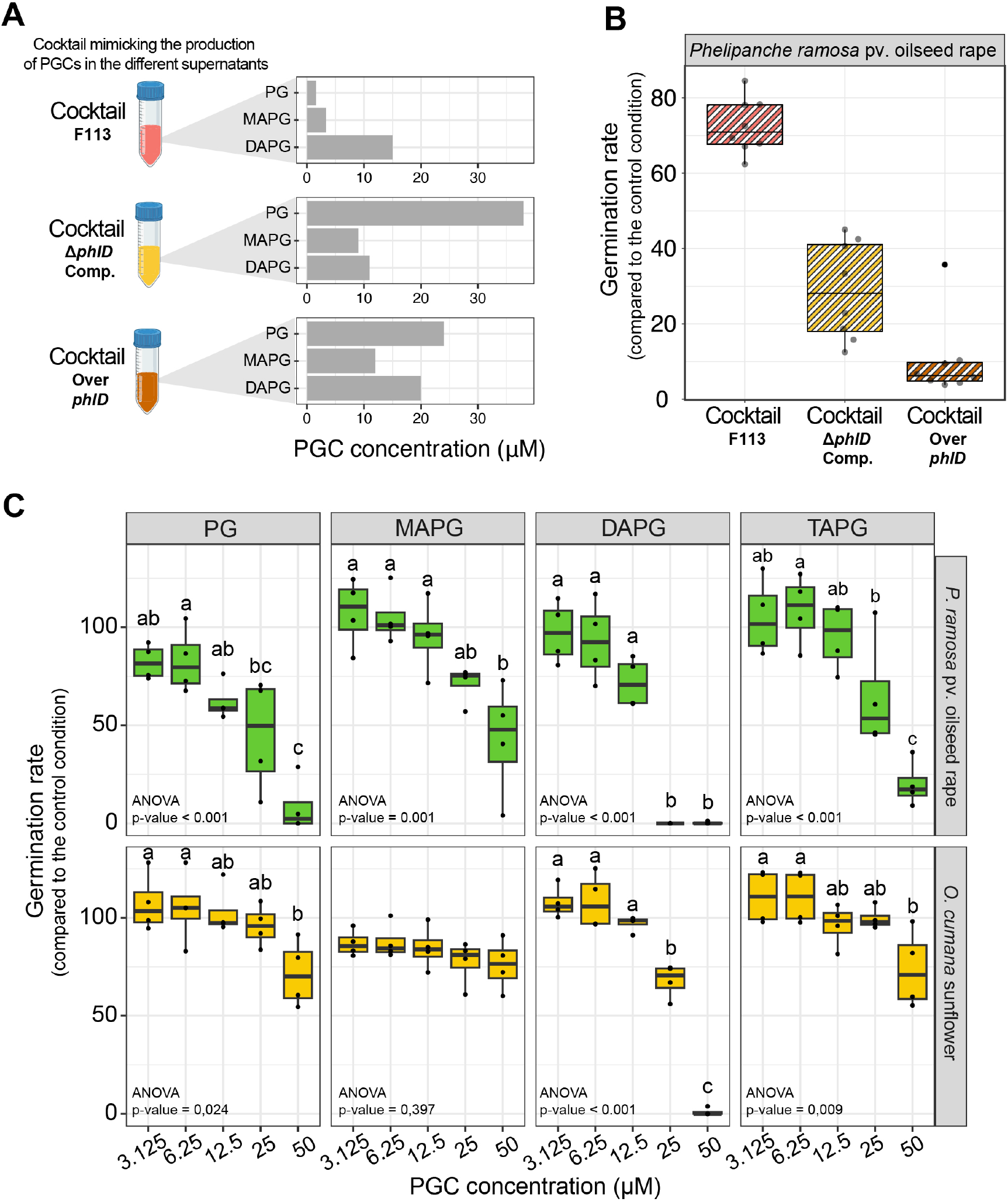
Individual contribution of the different PGCs to the inhibition of the germination of *P. ramosa* pv. oilseed rape and *O. cumana* sunflower. **A**: Quantification of the pure PGCs added into the different cocktails mimicking the concentration detected in the bacterial supernatants. PG: Phloroglucinol; MAPG: monoacetylphloroglucinol; DAPG: Diacetylphloroglucinol; TAPG: Triacetylphloroglucinol. **B**: Effect of the PGCs cocktails on the germination rate of *P. ramosa* pv oilseed rape. **C**: Impact of PG, MAPG, DAPG and TAPG on the germination capacity of different *P. ramosa* pv. oilseed rape and *O. cumana* sunflower *in vitro*. In all the conditions, broomrape seed germination was induced by adding the germination stimulant GR24 (1 μM). Results are expressed as a percentage of germination of the non-inoculated methanol control. Statistical differences were assessed by ANOVA and Kruskal-Wallis test using a Bonferroni correction and are indicated with letters. The horizontal lines indicate the interquartile range with the center representing the median. This experiment was repeated four times independently.

The same levels of inhibition were observed with the different molecular cocktails as in the experiment with bacterial supernatants. Indeed, the cocktails mimicking the concentration of PGCs in the supernatants of complemented and the overproducing strains most inhibited the germination of *P. ramosa* pv. oilseed rape (**Figure 2B**). Thus, as observed with complex supernatants, the inhibition of *P. ramosa* germination is dependent on the composition of the PGCs and their relative concentrations (**Figure 2A and 2B**).

Furthermore, we wanted to determine the individual contribution of these PGCs to the inhibition of broomrape germination by testing five different concentrations of each PGC (**Figure 2C**). First, we performed this inhibition assays on the four previously used broomrape species. However, the presence of 0.33 % of methanol in the PGC solutions inhibited seed germination of *P. ramosa* pv. tobacco and *P. ramosa* pv. hemp (Data not shown). Contrariwise to what was observed with the bacterial supernatants (**Supplementary Figure 1**), the germination rate of *P. ramosa* pv. oilseed rape was more affected following the exposure to PGCs than *O. cumana* sunflower (two-way ANOVA showed a broomrape effect, p-value < 0.001) (**Figure 2C**). Indeed, a reduction of more than 50 % of *O. cumana* germination was observed only where the seeds were exposed to 50 μM of DAPG. Remarkably, MAPG did not affect the germination rate of *O. cumana*. On the contrary, the germination rate of *P. ramosa* pv. oilseed rape started to be affected by the addition of PGCs at 12.5 μM. DAPG was the most effective PGC to disable broomrape germination with 100 % of inhibition at 25_μM and 50 μM for *P. ramosa* pv. oilseed rape and 100 % inhibition at 50 μM for *O. cumana* sunflower. Indeed, the estimated median lethal concentration (LC50) is the lowest for the DAPG for both broomrape species tested (20.0 μM for *P. ramosa* pv. oilseed rape and 29.1 μM for *O. cumana* (**Supplementary Figure 2**). At the end of the experiment, we recovered the seeds of *P. ramosa* treated with 50 μM of the different PGCs and washed those 3 times with phosphate buffer to remove PGC traces. Then we added again the GR24 germination stimulant. We observed that the inhibition by the PGC is irreversible since none of the seed treated with PGC germinated. Moreover, for all PGCs except for DAPG, between 25 μM and 50 μM, the inhibition effects were associated with a brown coloration of the seeds or the radicles (**Supplementary Figure 3**).

### The inoculation of PGC-producing *Pseudomonas* reduced the infection of *P. ramosa* on *Brassica napus*

As the DAPG showed the best inhibition of broomrape germination *in vitro*, we tested the application of DAPG (50 μM and 250 μM) or bacteria (F113 and Δ*phlD)* on *Brassica napus* to reduce the infection by *P. ramosa* pv. oilseed rape in greenhouse conditions. We evaluated the developmental stage of *P. ramosa* using the development scale available in the **Supplementary Figure 4**. We observed a significant reduction (47%) of the number of *P. ramosa* attached to the root of oilseed rape when F113 was inoculated (**Figure 3A**). This effect was not detected when the Δ*phlD* mutant was inoculated. Interestingly, at the end of the experiment, we observed a reduction of the proportion of early stage infections (stage 1 and 2; qualitative infection scale available in **Supplementary Figure 4**) in the condition inoculated with F113 or Δ*phlD* compared to the control (**Figure 3B**). On the contrary, the application of DAPG at 50 or 250 μM (v = 5mL) did not affect the infection of *P. ramosa* (**Figure 3A and 3B**). We also measured the effect of the bacterial inoculation and the application of DAPG on the physiology of *Brassica napus*. We observed that the biomass of the root system was significantly lower than the control condition when F113 was inoculated (**Figure 4**). Conversely, the shoot biomass was significantly higher than the control condition when the DAPG was added (**Figure 4**). We also looked at the nutrition capacity of *B. napus* using an elemental profiling approach (i.e., ionomic) to investigate potential switch of elements according to the treatments we applied. For all the tested conditions, we did not observe a main significant switch of the ion profile inside the shoots of *B. napus* (**Figure 5A and 5B**). However, the proportions of certain ions changed according to the treatment applied. On the one hand, the inoculation of F113 lead to a significant increase of sodium and a decrease of manganese quantity, whereas the mutant Δ*phlD* was associated to a decrease of potassium (P) (**Figure 5C**). On the other hand, the addition of DAPG was associated to a significant increase of antimony (Sb), beryllium (Be) and a reduction of nickel (Ni) in rapeseed shoots (**Figure 5D**).

**Figure 3:**
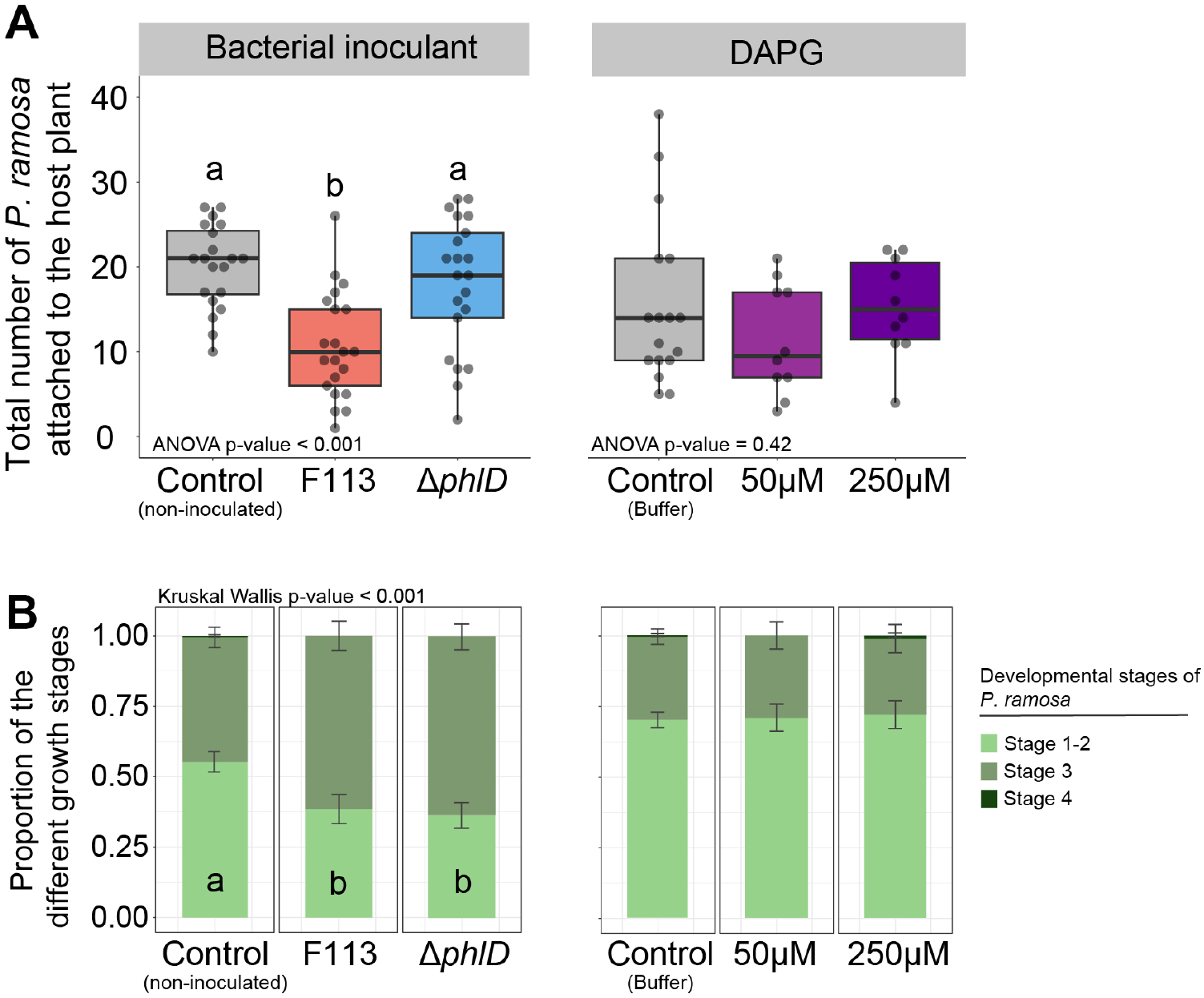
Impact of the inoculation of PGC-producing *Pseudomonas* and pure DAPG on the infection level by *P. ramosa* pv. oilseed rape on *Brassica napus* in greenhouse conditions. **A**: Evaluation of the number of attached *P. ramosa* on the root system of *Brassica napus* after 50 days in the greenhouse. Different treatments were applied: bacterial inoculants (F113 and Δ*phlD* impaired in the production of DAPG, approximately five mL per pot of solutions at 2.10^6^ bacteria.mL^-1^) and DAPG application (five mL of pure DAPG at 50μM or 250 μM). The control condition of the bacterial inoculant experiment corresponds to the application of 5 ml with MgSO_4_ 10 mM while in the DAPG experiment, the control corresponds to the application of 5 ml with phosphate buffer supplemented with 1.25% of methanol. This experiment was performed in a mixture containing natural soil artificially infested with approximatively 300 *P. ramosa* seeds per liter of soil. **B**: Proportion of broomrapes attached to the root of *B. napus* according to their developmental stage. The developmental stage of *P. ramosa* was estimated according to the developmental scale available in **Supplementary Figure 4**. Statistical differences are indicated with letters (ANOVA and Fisher’s LSD tests, p < 0.05). The horizontal lines indicate the interquartile range with the center representing the median.

**Figure 4:**
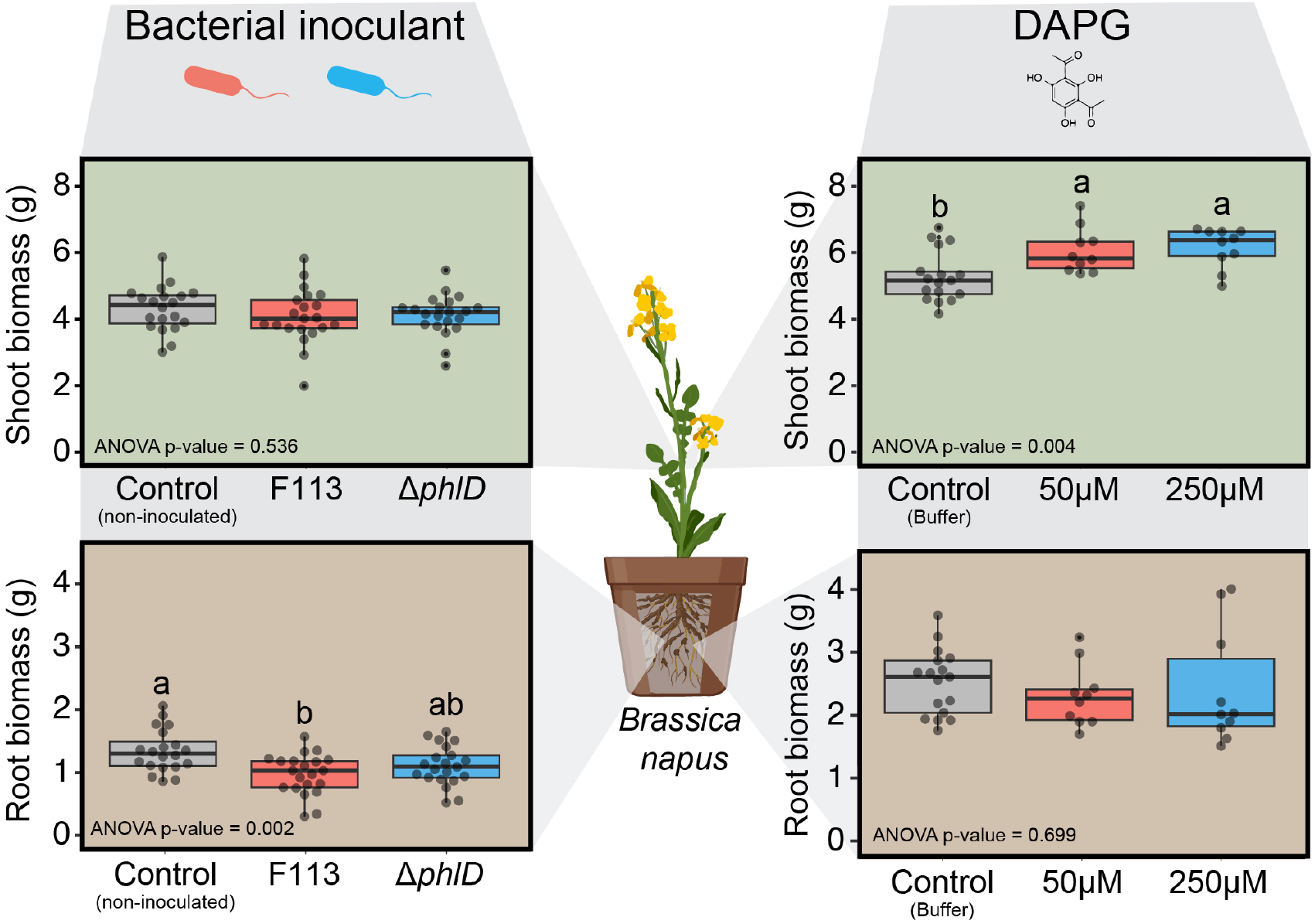
Effect of the inoculation of bacteria and the application of DAPG on the root and shoot biomasses of *Brassica napus*. We measured for all the tested conditions the shoot and root biomasses of *Brassica napus* 50 days after sowing. The control condition of the bacterial inoculant experiment corresponds to the application of 5 ml with MgSO_4_ 10 mM. In the DAPG experiment, the control corresponds to the application of 5 ml with phosphate buffer s supplemented with 1.25 % of methanol. The brown background of the boxplots corresponds to the root biomass data while the green background is associated with shoot biomass data. Statistical differences are indicated with letters (ANOVA and Fisher’s LSD tests, p < 0.05). The horizontal lines indicate the interquartile range with the center representing the median.

**Figure 5:**
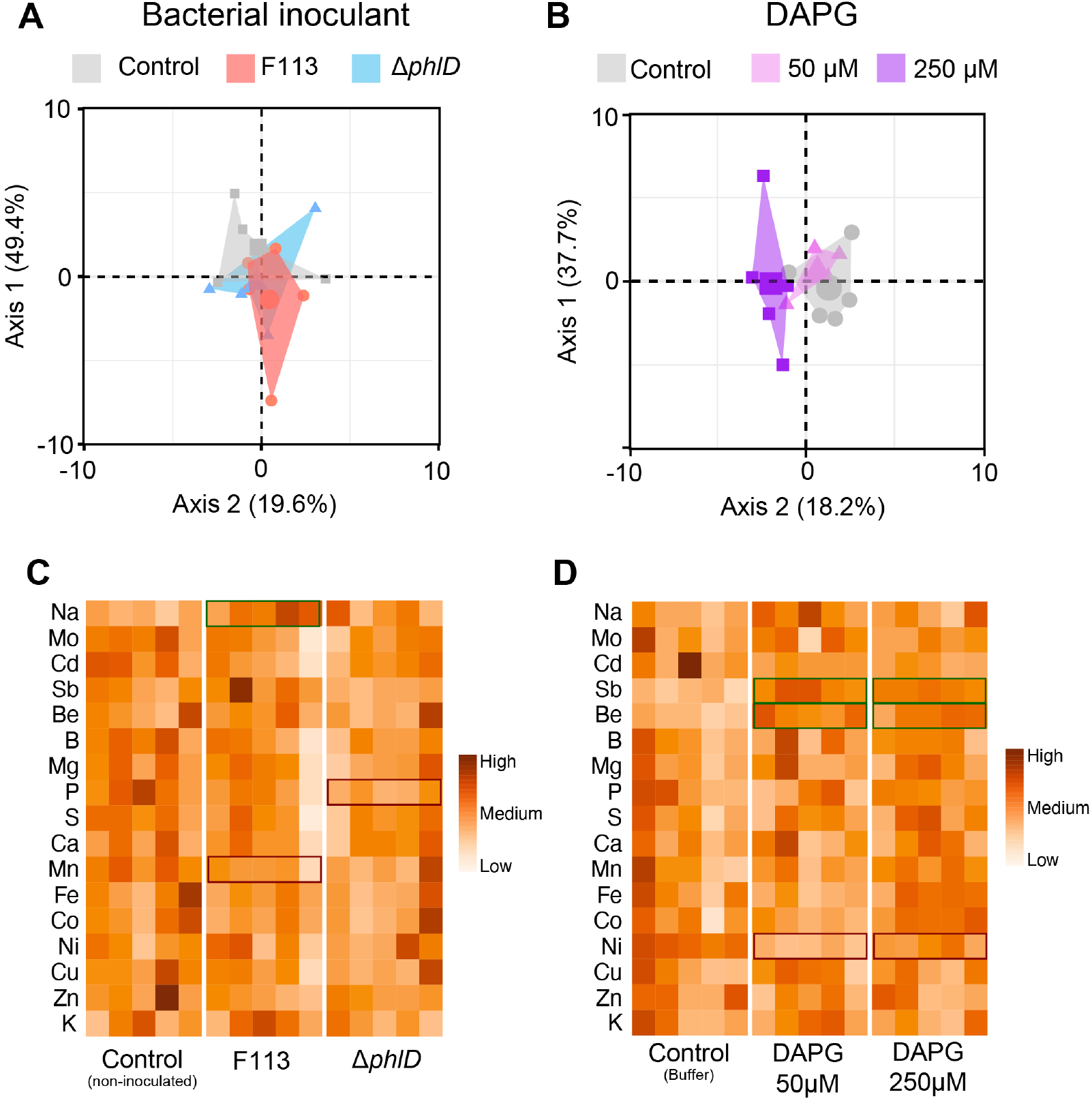
Impact of the bacterial inoculation and the application of DAPG on the ion profile of the shoot of *Brassica napus*. **A** and **B:** Principal component analysis of the element composition of the shoot of *Brassica napus* according to the different treatments applied (bacterial inoculants (**A**) and DAPG treatments (**B**)). **C** and **D** Heatmap showing the elemental profile of the different conditions. Significant differences between treatments and control were obtained with Student’s t-test. When Student’s t-test assumptions were not met, a Welch t-test was performed. *p-value < 0.05, **p-value < 0.01. The detailed data are available in **Supplementary Table 2** and **3**.

## Discussion

PGC-producing *Pseudomonas* are well described for their capacity to protect different crop plants from the infection by plant pathogens. Their biocontrol activity has been shown to be dependent on the production of antimicrobial compounds, including DAPG. In this study, we assessed whether the biocontrol activity of such bacterial strains can be extended to the protection of crops towards parasitic plants. We focused on the model environmental *Pseudomonas, P. ogarae* F113, which is known to display different plant-beneficial properties including the production of DAPG [33, 40]

Except for TAPG whose bacterial production remains to be demonstrated, F113 is able to synthesize, in different amounts, a cocktail of PGCs composed of PG, MAPG and DAPG (**Figure 1B**). It is worth noting that the production and the proportion of PGCs are variable among *Pseudomonas* strains [41] and depend on many environmental factors such as carbon sources [23], bacterial lifestyle (e.g., planktonic or biofilm) [42] or composition of plant exudates [43, 44]. The introduction of the *phlD* gene on a low-copy plasmid in the *phlD* mutant (Δ*phlD* Comp.) and in the wild-type F113 strain (Over *phlD)* modified the production of PGCs, these strains accumulating more PG and MAPG in their supernatants than F113. The production of different PGCs (as in the Over-*phlD* F113 derivative) leads to a stronger inhibitory effect on the germination of the different broomrapes tested (**Figure 1C**). We also confirmed these results using a cocktail of commercial PGCs in the same proportions (**Figure 2A and 2B**) and highlighted that each of the PGCs displayed a different inhibition capacity towards the germination of broomrapes (**Figure 2C**). Indeed, we showed that the DAPG provided the highest inhibition results. Islam and von Tiedemann (2011) tested the effect of DAPG and its derivatives on the zoosporogenesis and the motility of zoospores from *Plasmopara viticola* and *Aphanomyces cochlioides* [45]. They also observed that DAPG displayed the highest inhibitory effect compared to its derivatives. Similar results were observed on the mycelial growth of the plant pathogen *Pythium ultimum* [46]. Altogether, these results highlight the importance to consider the amount of the different precursors of DAPG produced by PGCs-producing *Pseudomonas* strains as they could also act as active compounds.

Although DAPG had been well described for its antimicrobial activity towards different kind of plant-pathogens (e.g., fungi, oomycetes) [22], its toxic effect on plants was also investigated on crop plants and appear to be associated to its concentration and the plant species [34]. A recent study analyzed the impact of DAPG at 50 μg.mL^-1^ (equivalent to 240 μM) on the germination rate of 69 wheat cultivars and showed strong differences according to cultivars [47]. This is in line with our results where the inhibition of germination was dependent on the concentration of DAPG and also different according to the broomrapes species and pathovar. Chae and colleagues (2020) identified several genes implicated in the sensitivity of *Arabidopsis thaliana* to exposure to DAPG [48]. These genes are involved in different metabolic pathways such as the tryptophan and monocarboxylic acid metabolisms and iron management [48]. The development of molecular tools to modify broomrape genomes would provide new insights on the molecular determinants responsible for their resistance and sensitivity towards PGCs.

The mode of action of DAPG and its derivatives at cellular level remains elusive. In our study, we observed a brown coloration of the broomrape radicles treated with PGCs (**Supplementary figure 3**). This coloration had previously been observed on tomato seedlings following the addition of 50 μM of DAPG [31]. At cellular levels, the sensitivity to DAPG was correlated with disruption of F-actin cytoskeleton in *A. cochlioides* hyphae [49] and with alterations of major physiological functions in yeast including the regulation of cellular responses to reactive oxygen stress and cell homeostasis [50]. Thus, the brown coloration of root tissues could be attributed to cell wall disorganization [49], impairment of mitochondrial functions [50, 51] and induction of oxidative burst [50]. Furthermore, in our experiment, the damage caused by the DAPG on broomrape seeds was irreversible since none of the broomrape seeds treated with DAPG were able to germinate after several washing steps and the addition of the GR24 germination inductor.

To determine if these compounds and PGCs-producing *Pseudomonas* strains are good candidates to prevent the infection of broomrapes, we performed greenhouse experiments with a natural soil which was infected with broomrape seeds. We used this soil to assess whether a bacterial inoculant or the pure DAPG can prevent the infection of oilseed rape by broomrapes. We observed that the wild type F113 was able to significantly reduce the number of broomrapes bound to *B. napus* roots compared to the mutant impaired in DAPG production. Interestingly, the number of broomrapes associated with new host infection (stage 1 and 2) in the condition where the bacteria were inoculated was significantly lower than in the control condition. Thus, the reduction of the infection is delayed and did not start immediately after the inoculation of the bacterial strains. This lag phase could be interpreted as the time needed for the bacterial inoculant to establish itself within the soil and/or root microbiome, and/or to produce DAPG in sufficient amounts. Bacteria can exert, as well, an indirect positive activity on oilseed rape leading to a decreased sensitivity to broomrape infection as cited previously by Cartry *et* al. [5]. Moreover, several reports showed that DAPG act as a signaling molecule inducing the plant systemic resistance [32] or the expression of its own biosynthetic genes [26]. Thus, the enrichment of the *B. napus* rhizo-microbiota with PGC-producing pseudomonads could stimulate other PGC-producing strains already present in the rhizosphere, leading to an increase of PGC concentrations in soil, thereby leading to detrimental effects on broomrape germination.

In addition, the inoculation of F113 leads to a decrease of the root system biomass of *B. napus*. It has been shown that the inoculation of PGC-producing *Pseudomonas* was linked to a decrease of the root length in *Arabidopsis thaliana* and other crop plants [31, 33, 52]. Thus, PGCs producing strains could limit the germination of broomrapes by two modes of action. The first one can be direct via the production of DAPG that inhibit the parasite seed germination whereas the second affects the root length of the host plant that *in fine* decreases the probability of contact between the host plant roots and broomrape seeds within the rhizosphere soil.

Conversely, the application of DAPG did not limit the infection of *P. ramosa* on *B. napus* roots. The DAPG applied may have not only specifically targeted the broomrape seeds but also interacted with the resident soil/root microbiomes as well as the oilseed plant. Thus, the amount of DAPG targeting the broomrape seeds might be insufficient to deliver a significant reduction of *P. ramosa* infection. Moreover, the soil chemical properties could impact the efficiency of DAPG inhibition. However, the application of the DAPG appeared to have an effect on the host plant by increasing oilseed rape shoot biomass (**Figure 4**). This gain of biomass was not associated in a significant shift in ion profile (**Figure 5**). Shoot accumulation of sodium (p-value < 0.05 for F113 treatment and < 0.1 for DAPG 50 μM treatment) appears to be linked to less parasitism. Sodium accumulation is generally considered as toxic for plants depending on plant species [53]. In our study, the accumulation of sodium does not affect the plant host but is correlated to a reduction of *P. ramosa* infection. Bacterial inoculation with *Pseudomonas* or *Azospirillum* was rather associated to a decrease of sodium in rapeseed leaf when exposed to harmful effects of salt stress [54]. Thus, improvement of sodium uptake in the host by DAPG-producing bacteria could be another factor associated to broomrape protection. As the nutrient flux from the host plant to the parasite is driven by an osmotic pressure differential between them [55], we hypothesize that the increase of sodium content in the host may reduce the level of nutrient uptake by the parasite and may impact its growth. Thus, sodium accumulation could be an interesting factor to study on oilseed rape cultivars in response to biocontrol agents.

Here, we showed that PGCs, and more specifically DAPG, combined different direct and indirect effects to protect oilseed rape against broomrape. DAPG and its derivatives could be interesting bioherbicides produced by *Pseudomonas* for preventing parasitic plant infestation as we did not observe any toxicity towards the host plant. The soil and root microbiome compositions were previously associated to natural suppressiveness towards parasitic weeds. Indeed, several studies claimed that the suppressiveness of *Orobanche* sp. as well as *Striga hermonthica* is associated with the presence of specific bacterial taxa, including *Pseudomonas* [11, 56]. Since, PGC-producing *Pseudomonas* can be followed in soil via qPCR approaches [57], determining the correlation between the community of PGC-producing *Pseudomonas* and the level of parasitic plant infection would bring new insights on the ecological role of these bacteria in suppressive soils.

## Conclusion

This study reinforces the interest of using microorganisms as natural solutions to regulate populations of pests and plant pathogens [58]. Indeed, this study is the first to discover and demonstrate the inhibitory effect of DAPG on parasitic plant germination, expanding the repertoire of plant-beneficial properties of environmental pseudomonads. Furthermore, we evidenced that PGCs produced by *P. ogarae* are the main and even the only compounds, that irreversibly inhibit the germination of broomrape species / pathovars tested. We obtained promising results during the experiments carried out in natural soil in greenhouse condition. Indeed, the inoculation of *P. ogarae* halved the infection of *P. ramosa* on its host plant, *Brassica napus*. The present work significantly expands our knowledge about the role that these plant-beneficial *Pseudomonas* play in the environment and provides new direction for the development of natural bioherbicides to ward off parasitic plant infection.

## Supporting information

Supplementary informations

## Author contributions

TL, JV, MR and CPC conceived the research and designed the experiment.

TL, MR, FG, SP, FWD and CPC performed the research.

TL, JV, FWD and CPC analyzed data.

TL, JV, FWD, CPC wrote the paper.

CPC found financial supports to this work.

## Acknowledgements

We thank the research team US2B of Nantes and more specifically Jean-Bernard Pouvreau for providing the broomrape seeds. We thank the company RAGT for providing seeds of the oilseed rape cultivars. We thank members of the Rhizo team of the Microbial Ecology unit at University Lyon 1 for their help in enumeration of broomrapes bound to *B. napus* roots in greenhouse. We are most grateful to PLATIN’ (Plateau d’Isotopie de Normandie) core facility for all element and isotope analysis used in this study. The platform ‘‘Serre’’ of FR BioEEnViS (University Lyon 1) was used to carry out this work.

## References

1. Parker C. Parasitic weeds: A world challenge. Weed Science 2012; 60: 269–276.

2. Bennett JR, Mathews S. Phylogeny of the parasitic plant family *Orobanchaceae* inferred from phytochrome A. Am J Bot 2006; 93: 1039–1051.

3. Joel DM. The new nomenclature of *Orobanche* and *Phelipanche*. Weed Research 2009; 49: 6–7.

4. Westwood JH. The physiology of the established parasite–host association. In: Joel DM, Gressel J, Musselman LJ (eds). Parasitic Orobanchaceae: Parasitic Mechanisms and Control Strategies. 2013. Springer, Berlin, Heidelberg, pp 87–114.

5. Cartry D, Steinberg C, Gibot-Leclerc S. Main drivers of broomrape regulation. A review. Agron Sustain Dev 2021; 41: 17.

6. Mutuku JM, Cui S, Yoshida S, Shirasu K. *Orobanchaceae* parasite–host interactions. New Phytolog 2021; 230: 46–59.

7. Aliche EB, Screpanti C, De Mesmaeker A, Munnik T, Bouwmeester HJ. Science and application of strigolactones. New Phytol 2020; 227: 1001–1011.

8. Haring SC, Flessner ML. Improving soil seed bank management. Pest Manag Sci 2018; 74: 2412–2418.

9. Rubiales D, Alcántara C, Pérez-de-Luque A, Gil J, Sillero J. Infection of chickpea *(Cicer arietinum*) by crenate broomrape (*Orobanche crenata*) as influenced by sowing date and weather conditions. dx.doi.org 2003; 23.

10. Huet S, Pouvreau J-B, Delage E, Delgrange S, Marais C, Bahut M, et al. Populations of the parasitic plant *Phelipanche ramosa* influence their seed microbiota. Front Plant Sci 2020; 11: 1075.

11. Kawa D, Thiombiano B, Shimels M, Taylor T, Walmsley A, Vahldick HE, et al. The soil microbiome reduces Striga infection of sorghum by modulation of host-derived signaling molecules and root development. 2022. bioRxiv.

12. Martinez L, Pouvreau J-B, Montiel G, Jestin C, Delavault P, Simier P, et al. Soil microbiota promotes early developmental stages of *Phelipanche ramosa* L. Pomel during plant parasitism on *Brassica napus* L. Plant Soil 2022.

13. Masteling R, Lombard L, de Boer W, Raaijmakers JM, Dini-Andreote F. Harnessing the microbiome to control plant parasitic weeds. Curr Opin Microbiol 2019; 49: 26–33.

14. Hasannejad S, Zad SJ, Alizade HM, Rahymian H. The effects of *Fusarium oxysporum* on broomrape *(Orobanche egyptiaca)* seed germination. Commun Agric Appl Biol Sci 2006; 71: 1295–1299.

15. Dadon T, Nun NB, Mayer AM. A factor from *Azospirillum brasilense* inhibits germination and radicle growth of *Orobanche aegyptiaca*. Isr J Plant Sci 2004; 52: 83–86.

16. Balthazar C, Joly DL, Filion M. Exploiting beneficial *Pseudomonas* spp. for cannabis production. Front Microbiol 2021; 12: 833172.

17. Silby MW, Winstanley C, Godfrey SAC, Levy SB, Jackson RW. *Pseudomonas* genomes: diverse and adaptable. FEMS Microbiol Rev 2011; 35: 652–680.

18. Haas D, Défago G. Biological control of soil-borne pathogens by fluorescent pseudomonads. Nat Rev Microbiol 2005; 3: 307–319.

19. Loper JE, Hassan KA, Mavrodi DV, Davis EW, Lim CK, Shaffer BT, et al. Comparative genomics of plant-associated *Pseudomonas* spp.: insights into diversity and inheritance of traits involved in multitrophic interactions. PLoS Genet 2012; 8: e1002784.

20. Almario J, Bruto M, Vacheron J, Prigent-Combaret C, Moёnne-Loccoz Y, Muller D. Distribution of 2,4-diacetylphloroglucinol biosynthetic genes among the *Pseudomonas* spp. reveals unexpected polyphyletism. Front Microbiol 2017; 8: 1218.

21. Achkar J, Xian M, Zhao H, Frost JW. Biosynthesis of phloroglucinol. J Am Chem Soc 2005; 127: 5332–5333.

22. Biessy A, Filion M. Phloroglucinol derivatives in plant-beneficial *Pseudomonas* spp.: biosynthesis, regulation, and functions. Metabolites 2021; 11: 182.

23. Shanahan P, O’sullivan DJ, Simpson P, Glennon JD, O’gara F. Isolation of 2,4-diacetylphloroglucinol from a fluorescent pseudomonad and investigation of physiological parameters influencing its production. Appl Environ Microbiol 1992; 58: 353–358.

24. Yu X-Q, Yan X, Zhang M-Y, Zhang L-Q, He Y-X. Flavonoids repress the production of antifungal 2,4-DAPG but potentially facilitate root colonization of the rhizobacterium *Pseudomonas fluorescens*. Environ Microbiol 2020; 22: 5073–5089.

25. Brodhagen M, Henkels MD, Loper JE. Positive autoregulation and signaling properties of pyoluteorin, an antibiotic produced by the biological control organism *Pseudomonas fluorescens* Pf-5. Appl Environ Microbiol 2004; 70: 1758–1766.

26. Maurhofer M, Baehler E, Notz R, Martinez V, Keel C. Cross talk between 2,4-diacetylphloroglucinol-producing biocontrol pseudomonads on wheat roots. Appl Environ Microbiol 2004; 70: 1990–1998.

27. Combes-Meynet E, Pothier JF, Moënne-Loccoz Y, Prigent-Combaret C. The *Pseudomonas* secondary metabolite 2,4-diacetylphloroglucinol is a signal inducing rhizoplane expression of *Azospirillum* genes involved in plant-growth promotion. Mol Plant Microbe Interact 2011; 24: 271–284.

28. Iavicoli A, Boutet E, Buchala A, Métraux J-P. Induced systemic resistance in *Arabidopsis thaliana* in response to root inoculation with *Pseudomonas fluorescens* CHA0. Mol Plant Microbe Interact 2003; 16: 851–858.

29. Bakker PAHM, Pieterse CMJ, van Loon LC. Induced systemic resistance by fluorescent *Pseudomonas* spp. Phytopathology 2007; 97: 239–243.

30. Phillips DA, Fox TC, King MD, Bhuvaneswari TV, Teuber LR. Microbial products trigger amino acid exudation from plant roots. Plant Physiol 2004; 136: 2887–2894.

31. Brazelton JN, Pfeufer EE, Sweat TA, Gardener BBM, Coenen C. 2,4-Diacetylphloroglucinol alters plant root development. Mol Plant Microbe Interact 2008; 21: 1349–1358.

32. Weller DM, Mavrodi DV, van Pelt JA, Pieterse CMJ, van Loon LC, Bakker PAHM. Induced systemic resistance in *Arabidopsis thaliana* against *Pseudomonas syringae* pv. tomato by 2,4-diacetylphloroglucinol-producing *Pseudomonas fluorescens*. Phytopathology 2012; 102: 403–412.

33. Vacheron J, Desbrosses G, Renoud S, Padilla R, Walker V, Muller D, et al. Differential contribution of plant-beneficial functions from *Pseudomonas kilonensis* F113 to root system architecture alterations in *Arabidopsis thaliana* and *Zea mays*. Mol Plant Microbe Interact 2018; 31: 212–223.

34. Keel C. Suppression of root diseases by *Pseudomonas fluorescens* CHA0: Importance of the bacterial secondary metabolite 2,4-diacetylphloroglucinol. Mol Plant Microbe Interact 1992; 5: 4.

35. Khan F, Tabassum N, Bamunuarachchi NI, Kim Y-M. Phloroglucinol and its derivatives: antimicrobial properties toward microbial pathogens. J Agric Food Chem 2022; 70: 4817–4838.

36. King EO, Ward MK, Raney DE. Two simple media for the demonstration of pyocyanin and fluorescin. J Lab Clin Med 1954; 44: 301–307.

37. Pouvreau J-B, Gaudin Z, Auger B, Lechat M-M, Gauthier M, Delavault P, et al. A high-throughput seed germination assay for root parasitic plants. Plant Methods 2013; 9: 32.

38. El Zemrany H, Cortet J, Peter Lutz M, Chabert A, Baudoin E, Haurat J, et al. Field survival of the phytostimulator *Azospirillum lipoferum* CRT1 and functional impact on maize crop, biodegradation of crop residues, and soil faunal indicators in a context of decreasing nitrogen fertilisation. Soil Biol Biochem 2006; 38: 1712–1726.

39. Lurthy T, Cantat C, Jeudy C, Declerck P, Gallardo K, Barraud C, et al. Impact of bacterial siderophores on iron status and ionome in pea. Front Plant Sci 2020; 11: 730.

40. Redondo-Nieto M, Barret M, Morrisey JP, Germaine K, Martínez-Granero F, Barahona E, et al. Genome sequence of the biocontrol strain *Pseudomonas fluorescens* F113. J Bacteriol 2012; 194: 1273–1274.

41. Duffy BK, Défago G. Environmental factors modulating antibiotic and siderophore biosynthesis by *Pseudomonas fluorescens* biocontrol strains. Appl Environ Microbiol 1999; 65: 2429–2438.

42. Rieusset L, Rey M, Muller D, Vacheron J, Gerin F, Dubost A, et al. Secondary metabolites from plant-associated *Pseudomonas* are overproduced in biofilm. Microb Biotechnol 2020; 13: 1562–1580.

43. de Werra P, Huser A, Tabacchi R, Keel C, Maurhofer M. Plant-and microbe-derived compounds affect the expression of genes encoding antifungal compounds in a pseudomonad with biocontrol activity. Appl Environ Microbiol 2011; 77: 2807–2812.

44. Rieusset L, Rey M, Gerin F, Wisniewski-Dyé F, Prigent-Combaret C, Comte G. A cross-metabolomic approach shows that wheat interferes with fluorescent *Pseudomonas* physiology through its root metabolites. Metabolites 2021; 11: 84.

45. Islam MT, von Tiedemann A. 2,4-Diacetylphloroglucinol suppresses zoosporogenesis and impairs motility of Peronosporomycete zoospores. World J Microbiol Biotechnol 2011; 27: 2071–2079.

46. de Souza JT, Arnould C, Deulvot C, Lemanceau P, Gianinazzi-Pearson V, Raaijmakers JM. Effect of 2,4-diacetylphloroglucinol on *pythium:* cellular responses and variation in sensitivity among propagules and species. Phytopathology 2003; 93: 966–975.

47. Yang M, Thomashow LS, Weller DM. Evaluation of the phytotoxicity of 2,4-diacetylphloroglucinol and *Pseudomonas brassicacearum* Q8r1-96 on different wheat cultivars. Phytopathology 2021; 111: 1935–1941.

48. Chae D-H, Kim D-R, Cho G, Moon S, Kwak Y-S. Genome-wide investigation of 2,4-diacetylphloroglucinol protection genes in *Arabidopsis thaliana*. Mol Plant-Microbe Interact 2020; 33: 1072–1079.

49. Islam MT, Fukushi Y. Growth inhibition and excessive branching in *Aphanomyces cochlioides* induced by 2,4-diacetylphloroglucinol is linked to disruption of filamentous actin cytoskeleton in the hyphae. World J Microbiol Biotechnol 2010; 26: 1163–1170.

50. Kwak Y-S, Han S, Thomashow LS, Rice JT, Paulitz TC, Kim D, et al. *Saccharomyces cerevisiae* genome-wide mutant screen for sensitivity to 2,4-diacetylphloroglucinol, an antibiotic produced by *Pseudomonas fluorescens*. Appl Environ Microbiol 2011; 77: 1770–1776.

51. Troppens DM, Dmitriev RI, Papkovsky DB, O’Gara F, Morrissey JP. Genome-wide investigation of cellular targets and mode of action of the antifungal bacterial metabolite 2,4-diacetylphloroglucinol in *Saccharomyces cerevisiae*. FEMS Yeast Res 2013; 13: 322–334.

52. De Leij FA, Dixon-Hardy JE, Lynch JM. Effect of 2,4-diacetylphloroglucinol-producing and non-producing strains of *Pseudomonas fluorescens* on root development of pea seedlings in three different soil types and its effect on nodulation by *Rhizobium*. Biol Fertil Soils 2002; 35: 114–121.

53. Kronzucker HJ, Coskun D, Schulze LM, Wong JR, Britto DT. Sodium as nutrient and toxicant. Plant Soil 2013; 369: 1–23.

54. Farhangi-Abriz S, Tavasolee A, Ghassemi-Golezani K, Torabian S, Monirifar H, Rahmani HA. Growth-promoting bacteria and natural regulators mitigate salt toxicity and improve rapeseed plant performance. Protoplasma 2020; 257: 1035–1047.

55. Shen H, Ye W, Hong L, Huang H, Wang Z, Deng X, et al. Progress in parasitic plant biology: host selection and nutrient transfer. Plant Biol (Stuttg) 2006; 8: 175–185.

56. Zermane N, Souissi T, Kroschel J, Sikora R. Biocontrol of broomrape (*Orobanche crenata* Forsk. and *Orobanche foetida* Poir.) by *Pseudomonas fluorescens* isolate Bf7-9 from the faba bean rhizosphere. Biocontrol Sci Technol 2007; 17: 483–497.

57. Almario J, Moënne-Loccoz Y, Muller D. Monitoring of the relation between 2,4-diacetylphloroglucinol-producing *Pseudomonas* and *Thielaviopsis basicola* populations by real-time PCR in tobacco black root-rot suppressive and conducive soils. Soil Biol Biochem 2013; 57: 144–155.

58. Vurro M. Are root parasitic broomrapes still a good target for bioherbicide control? Pest Manag Sci 2023.

