## Supplementary informations for "Inhibition of broomrape germination by 2,4-diacetylphloroglucinol produced by environmental *Pseudomonas*"

#### Supplementary Figures

**Supplementary Figure 1:** The sensitivity of broomrapes to bacterial supernatants is species and pathovar dependent.

**Supplementary Figure 2:** Dose-response of two different broomrape species following their exposure to different PGCs *in vitro*.

**Supplementary Figure 3:** The exposure of *P. ramosa* seeds to PGCs (PG, MAPG, DAPG or TAPG) and PGCs-producing *Pseudomonas* strains supernatant leads to the apparition of a brown coloration.

**Supplementary Figure 4:** Infectivity scale of *Phelipanche ramosa* during the greenhouse experiments.

#### Supplementary Tables

**Supplementary Table 1:** List of the bacterial strains used in this study.

**Supplementary Table 2:** Impact of the inoculation of F113 or Δ*phlD* on the elemental composition of the leaves of *B. napus*. (µg.g$^{-1}$ of dry biomass, except for Mg, P, S, Ca, Fe and K i.e., mg.g$^{-1}$ of dry biomass).

**Supplementary Table 3:** Impact of the application of two concentrations of DAPG on the elemental composition of the leaves of *B. napus*. (µg.g$^{-1}$ of dry biomass, except for Mg, P, S, Ca, Fe and K i.e., mg.g$^{-1}$ of dry biomass).

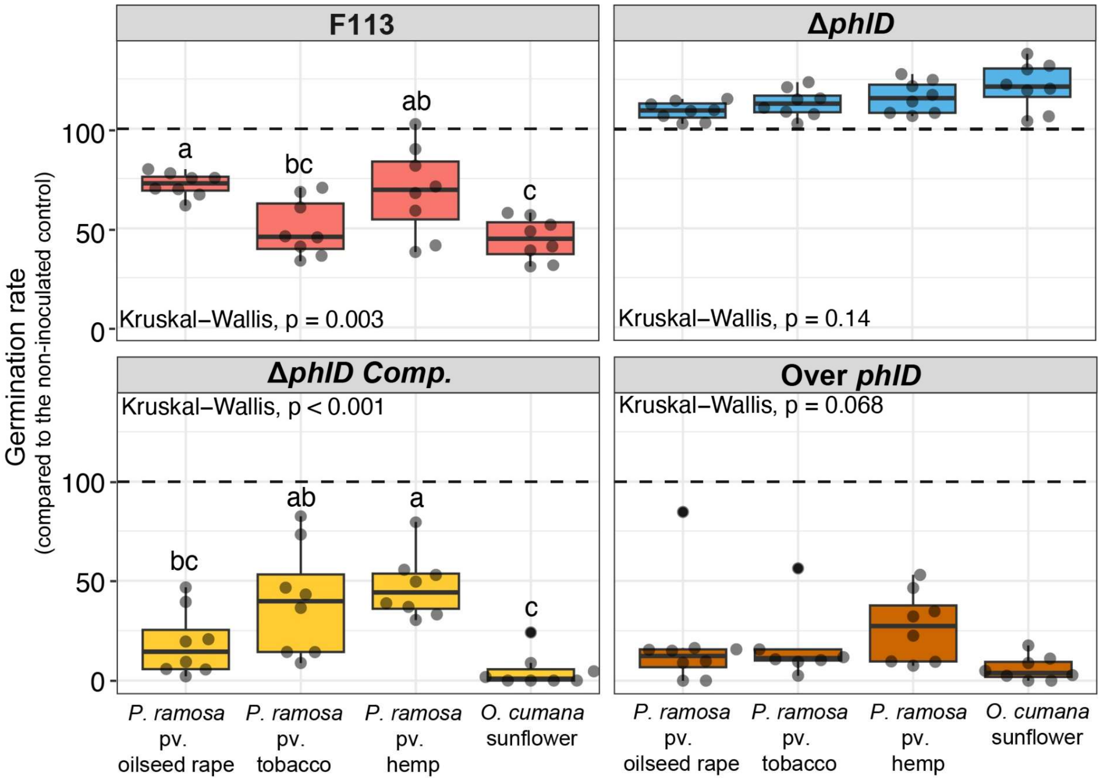


**Supplementary Figure 1: The sensitivity of broomrapes to bacterial supernatants is species and pathovar dependent.** Impact of supernatants from F113 and its derivatives on the germination capacity of different broomrapes *in vitro*. The supernatants as well as the control condition were supplemented with 1 µM of the germination stimulant (GR24). Results are expressed as percentage of germination of the non-inoculated control. Statistical differences were assessed by ANOVA and Kruskal-Wallis test using a Bonferroni correction and are indicated with letters. The horizontal lines indicate the interquartile range with the center representing the median.

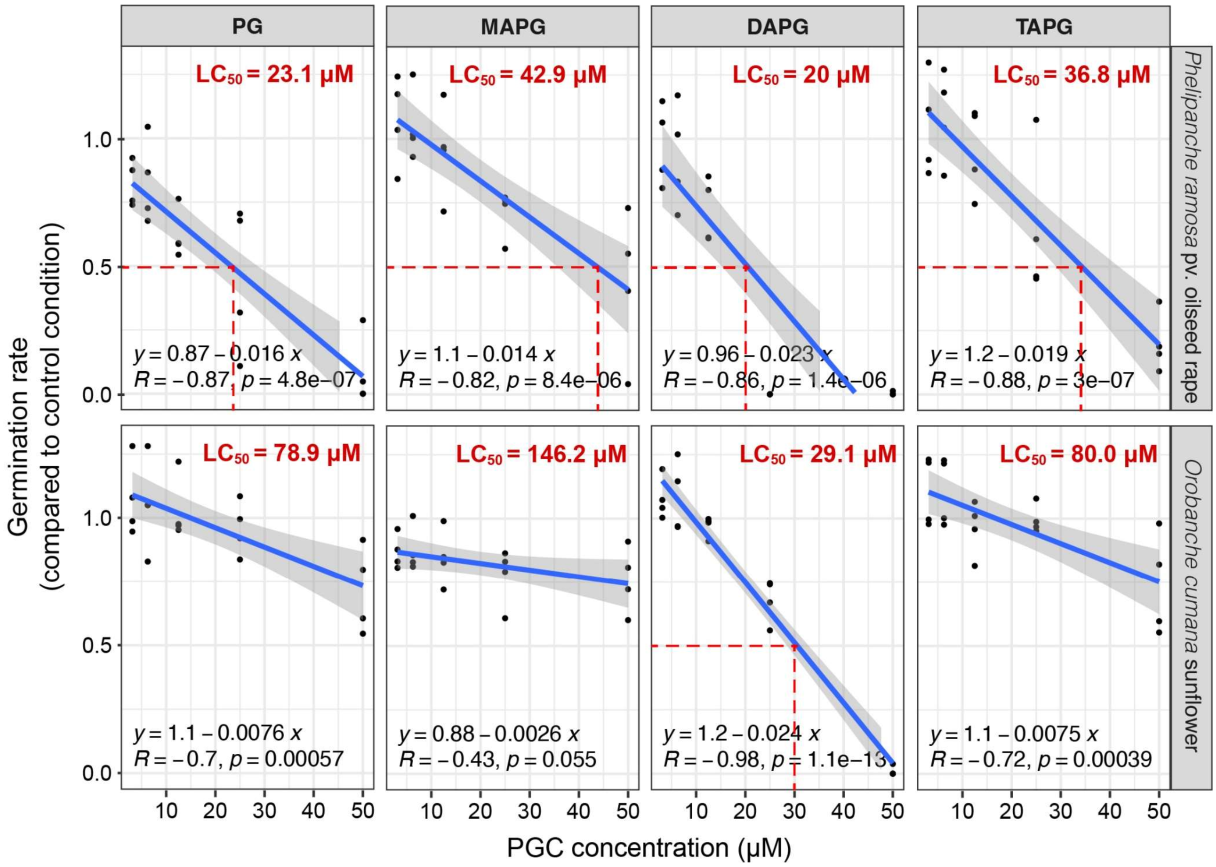


**Supplementary Figure 2: Dose-response of two different broomrape species following their exposure to different PGCs *in vitro*.** The LC50 (Lethal concentration 50) was estimated thanks to a linear regression model. Correlation coefficient were calculated using Spearman correlation test. PG: Phloroglucinol; MAPG: Monoacetylphloroglucinol; DAPG: Diacetylphloroglucinol; TAPG: Triacetylphloroglucinol.

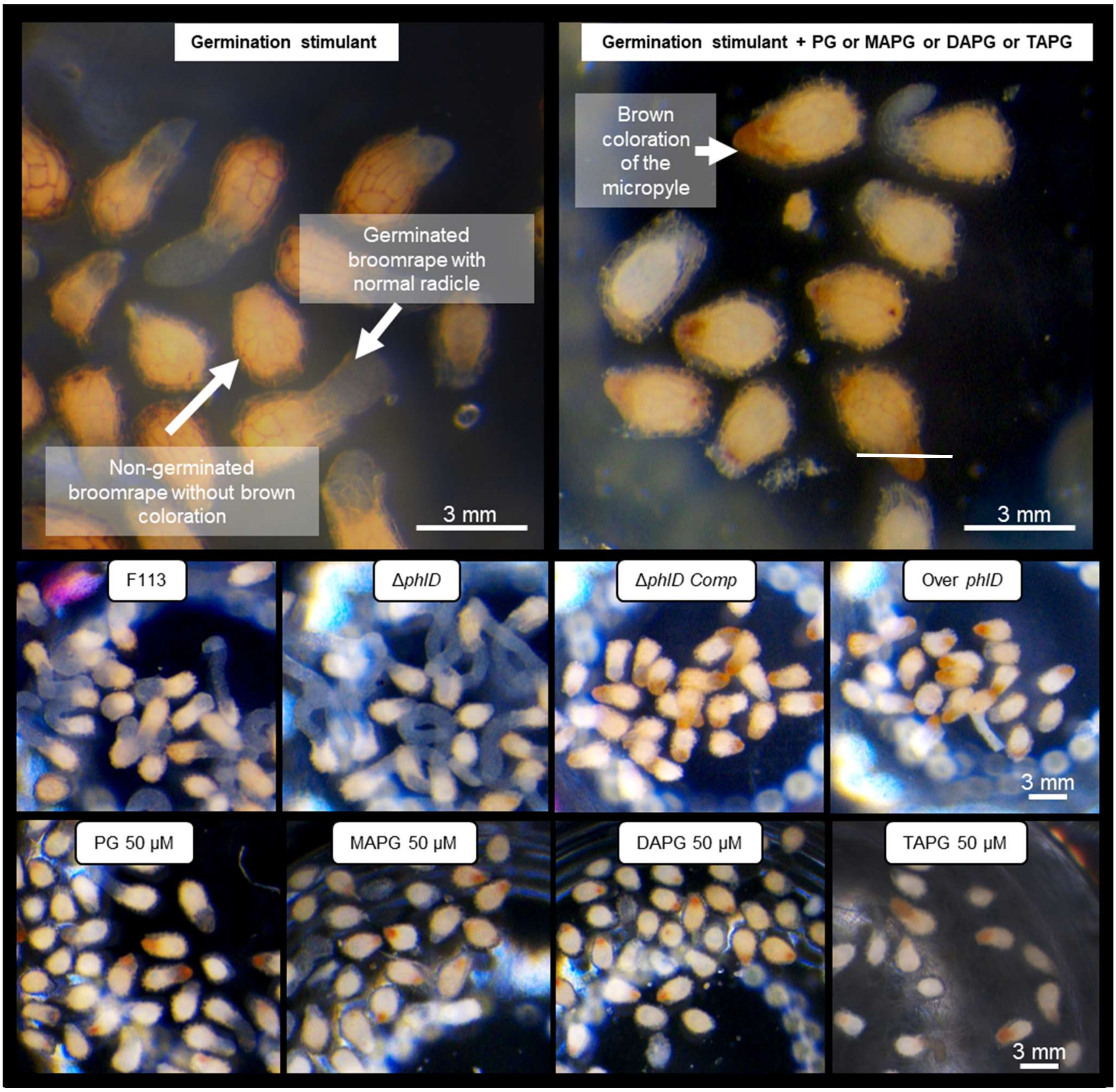


**Supplementary Figure 3: The exposure of *P. ramosa* seeds to PGCs (PG, MAPG, DAPG or TAPG) and PGCs-producing *Pseudomonas* strains supernatant leads to the apparition of a brown coloration.** Seeds were photographed using camera (AxioCam MRc5) attached to binocular loupe, zoom x20.

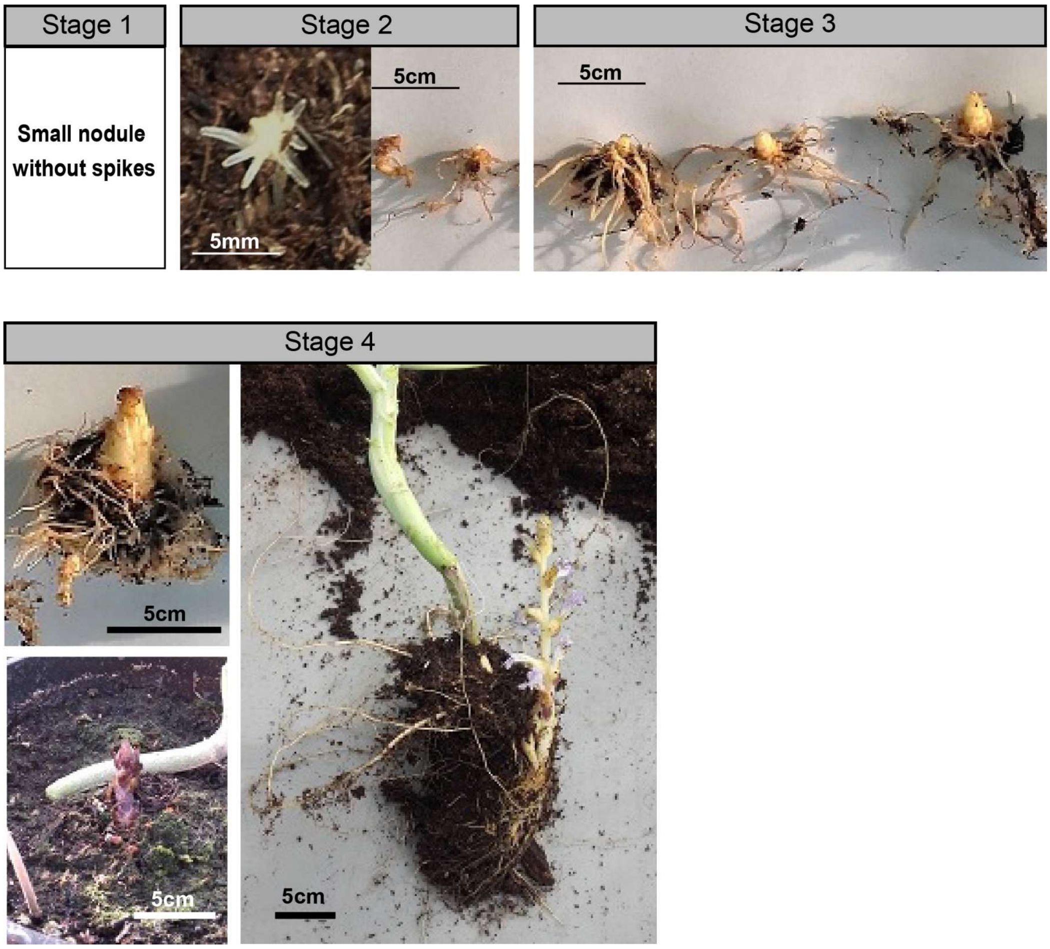


**Supplementary Figure 4: Infectivity scale of *Phelipanche ramosa* used during the greenhouse experiments.** *Stage 1* corresponds to small tubercules without spikes (<2 mm of diameter); *Stage 2* corresponds to nodule with spikes without visible broomrape bud; *Stage 3* corresponds to the apparition of broomrape bud; *Stage 4* encompasses the development of floral stem and its emergence outside the soil.

**Supplementary Table 1:** List of the bacterial strains used in this study.

| Strain names | Genotype or relevant characteristics[1] | Reference or source |
|---|---|---|
| F113 | *Pseudomonas ogarae* F113 wild type | [1] |
| Δ*phlD* | Deletion mutant of *phlD*, impaired in the production of DAPG. | [2] |
| Δ*phlD Comp* | Δ*phlD* containing the low-copy plasmid pBBR1-MCS5:*phlD*, restoration of the DAPG production. $Gm^R$ | [2] |
| Over *phlD* | F113 wild type containing the low-copy plasmid pBBR1-MCS5:*phlD*, enhancement of the DAPG production. $Gm^R$ | This work |

[1] $Gm^R$, gentamicin resistance.

**Supporting information Table 2:** Impact of the inoculation of F113 or Δ*phlD* on the elemental composition of the leaves of *B. napus*. (μg.g$^{-1}$ of dry biomass, except for Mg, P, S, Ca, Fe and K i.e., mg.g$^{-1}$ of dry biomass).

| | Control | | F113 | | Δ*phlD* | |
|---|---|---|---|---|---|---|
| | **Mean** | *Average deviation* | **Mean** | *Average deviation* | **Mean** | *Average deviation* |
| **Shoot dry weight (g)** | 4.3 | *0.2* | 4.1 | *0.2* | 4.1 | *0.1* |
| **Number of attached *P. ramosa*** | 20.1 | *1.1* | 10.7 | *1.4* | 18 | *1.7* |
| **Na** | 3923.89 | *374.92* | 5545.03* | *700.82* | 4610.88 | *832.7* |
| **K** | 125.74 | *12.90* | 132.62 | *15.49* | 120.73 | *10.11* |
| **Mo** | 78.99 | *9.72* | 60.55 | *17.08* | 66.32 | *11.92* |
| **Cd** | 1.14 | *0.19* | 0.84 | *0.19* | 1.02 | *0.11* |
| **Sb** | 0.10 | *0.03* | 0.15 | *0.057* | 0.06 | *0.013* |
| **Be** | 0.07 | *0.02* | 0.07 | *0.01* | 0.07 | *0.02* |
| **B** | 158.28 | *15.78* | 133.64 | *24.23* | 137.89 | *11.96* |
| **Mg** | 39.81 | *2.57* | 38.16 | *3.4* | 37.34 | *2.73* |
| **P** | 27.17 | *2.33* | 23.65 | *2.35* | 23.32* | *1.12* |
| **S** | 86.98 | *4.82* | 79.31 | *9.14* | 82.34 | *5.43* |
| **Ca** | 188.25 | *15.50* | 180.74 | *13.95* | 187.01 | *12.39* |
| **Mn** | 220.51 | *12.55* | 192.44* | *14.87* | 201.84 | *20.94* |
| **Fe** | 1.53 | *0.60* | 1.16 | *0.25* | 1.08 | *0.52* |
| **Co** | 1.03 | *0.29* | 0.8 | *0.15* | 0.86 | *0.36* |
| **Ni** | 3.00 | *0.49* | 3.06 | *0.96* | 3.17 | *0.74* |
| **Cu** | 30.29 | *1.94* | 28.1 | *2.37* | 28.57 | *2.57* |
| **Zn** | 192.14 | *50.90* | 146.92 | *21.29* | 165.45 | *10.59* |

Sodium (Na), Molybdenum (Mo), Cadmium (Cd), Antimony (Sb), Beryllium (Be), Bore (B), Magnesium (Mg), Phosphorus (P), Sulphur (S), Calcium (Ca), Manganese (Mn), Iron (Fe), Cobalt (Co), Nickel (Ni), Copper (Cu), Zinc (Zn), Potassium (K). n=5 samples of pooled of four plants. Significant differences between treatments and control were assessed using either t-tests or Welch t-test when Student's t-test assumptions were not met. *, p<0.05; **, p<0.01. Red and green percentages correspond to significant negative and positive impact on ionome respectively.

**Supporting information Table 3:** Impact of the application of two concentration of DAPG on the elemental composition of the leaves of *B. napus*. (µg.g$^{-1}$ of dry biomass, except for Mg, P, S, Ca, Fe and K i.e., mg.g$^{-1}$ of dry biomass).

| | Control | | DAPG 50 µM | | DAPG 250 µM | |
|---|---|---|---|---|---|---|
| | **Mean** | *Average deviation* | **Mean** | *Average deviation* | **Mean** | *Average deviation* |
| **Shoot dry weight (g)** | 5.3 | *0.18* | 6.15 | *0.19* | 6.1 | *0.21* |
| **Number of attached *P. ramosa*** | 15.4 | *2.4* | 11.4 | *2.1* | 15.3 | *1.9* |
| **Na** | 320.28 | *24.65* | 411.35• | *62.92* | 353.63 | *59.27* |
| **K** | 66.87 | *10.01* | 68.43 | *7.12* | 63.46 | *6.44* |
| **Mo** | 48.67 | *4.88* | 49.49 | *4.16* | 47.78 | *1.45* |
| **Cd** | 0.84 | *0.43* | 0.57 | *0.02* | 0.54 | *0.05* |
| **Sb** | 0.03 | *0.01* | 0.12* | *0.05* | 0.10*** | *0.01* |
| **Be** | 0.052 | *0.002* | 0.063** | *0.004* | 0.063** | *0.003* |
| **B** | 94.23 | *9.74* | 97.37 | *13.99* | 94.95 | *5.81* |
| **Mg** | 18.06 | *1.74* | 18.71 | *2.04* | 18.96 | *1.93* |
| **P** | 16.92 | *2.52* | 15.91 | *2.06* | 16.55 | *0.57* |
| **S** | 24.24 | *2.2* | 24.54 | *1570.8* | 25.7 | *2.23* |
| **Ca** | 99.85 | *11.15* | 103.94 | *16.9* | 103.81 | *13.21* |
| **Mn** | 152.49 | *17.05* | 149.76 | *7.74* | 158.29 | *5.6* |
| **Fe** | 349.61 | *60.9* | 286.13 | *21.79* | 373.13 | *34.86* |
| **Co** | 0.45 | *0.07* | 0.42 | *0.03* | 0.52 | *0.03* |
| **Ni** | 1.79 | *0.47* | 0.34* | *0.07* | 0.76* | *0.27* |
| **Cu** | 12.44 | *1.56* | 12.96 | *1.25* | 12.51 | *1.01* |
| **Zn** | 84.42 | *8.94* | 77.3 | *6.73* | 79.39 | *9.34* |

Sodium (Na), Molybdenum (Mo), Cadmium (Cd), Antimony (Sb), Beryllium (Be), Bore (B), Magnesium (Mg), Phosphorus (P), Sulphur (S), Calcium (Ca), Manganese (Mn), Iron (Fe), Cobalt (Co), Nickel (Ni), Copper (Cu), Zinc (Zn), Potassium (K). n=5 samples of pooled of two plants. Significant differences between treatments and control were assessed using either t-tests or Welch t-test when Student's t-test assumptions were not met. •, p<0.1 *, p<0.05; **, p<0.01. Red and green percentages correspond to significant negative and positive impact on ionome respectively.

### References

1. Shanahan P, O'sullivan DJ, Simpson P, Glennon JD, O'gara F. Isolation of 2,4-diacetylphloroglucinol from a fluorescent pseudomonad and investigation of physiological parameters influencing its production. *Appl Environ Microbiol* 1992; **58**: 353–358.

2. Vacheron J, Desbrosses G, Renoud S, Padilla R, Walker V, Muller D, et al. Differential contribution of plant-beneficial functions from *Pseudomonas kilonensis* F113 to root system architecture alterations in *Arabidopsis thaliana* and *Zea mays*. *MPMI* 2018; **31**: 212–223.
